# Accelerated electron transfer in nanostructured electrodes improves the sensitivity of electrochemical biosensors

**DOI:** 10.1101/2021.04.13.439686

**Authors:** Kaiyu Fu, Ji-Won Seo, Vladimir Kesler, Nicolo Maganzini, Brandon D. Wilson, Michael Eisenstein, H. Tom Soh

**Author notes:** These authors contributed equally to this work (Kaiyu Fu, Ji-Won Seo, Vladimir Kesler).

## Abstract

Electrochemical biosensors hold the exciting potential to integrate molecular detection with signal processing and wireless communication in a miniaturized, low-cost system. However, as electrochemical biosensors are miniaturized to the micron scale, their detection sensitivity degrades precipitously, thereby greatly reducing their utility in the context of molecular diagnostic applications. Studies have reported that nanostructured electrodes can greatly improve electrochemical biosensor sensitivity, but the underlying mechanism remains poorly understood, thus making it difficult to fully exploit this phenomenon to improve biosensor performance. In this work, we propose and experimentally validate a novel mechanism in which electron transfer is physically accelerated within nanostructured electrodes due to reduced charge screening, resulting in enhanced sensitivity. We show that this mechanism can be exploited to achieve up to 24-fold increase in signal and nearly four-fold lower limit-of-detection relative conventional planar electrodes. This accelerated electron transfer mechanism should prove broadly applicable for improving the performance of electrochemical biosensors.

## INTRODUCTION

Electrochemical biosensors have gained great interest in the past decade because they can be incorporated directly into very large-scale integrated circuits (VLSI).^1, 2^ This provides the exciting potential to fully couple biomolecular sensing with computation and communication in a miniaturized, low-cost system.^3-8^ Sensitivity is a key consideration for many biomedical applications, because many clinical biomarkers are present at nanomolar to picomolar concentrations, and the biosensor must achieve sufficient sensitivity in a complex background of interferent molecules.^9-14^ Unfortunately, the signal-to-noise ratio (SNR) of conventional electrochemical biosensors degrades precipitously when they are miniaturized to the micron scale,^15, 16^ reducing their sensitivity and making meaningful measurements of analyte concentrations challenging or even impossible in many cases.^17-19^

There have been a number of advances in the fabrication of nanostructured electrodes over the last decade,^20-25^ which achieve improved sensing properties relative to standard planar electrodes, such as increased signal levels and faster diffusion of redox species. In a seminal study, Kelley and co-workers demonstrated that nanostructured electrodes with high surface curvature, which they termed “nanoflowers”, can greatly enhance DNA detection compared to planar electrodes, with limits of detection (LOD) in the femtomolar range.^15^ Seker and co-workers have shown that similar improvements in sensitivity can also be achieved with nanoporous electrodes, with the additional benefit that the sensitivity and dissociation constant (*K*_*D*_) of the resulting sensors can be tuned by changing the size of the nanopores.^26, 27^ While the majority of prior work with nanostructured electrodes has been limited to the hybridization-based detection of nucleic acids, the use of electrode-coupled aptamers as a molecular recognition element can extend the same detection strategy to small molecules, peptides, and proteins. Indeed, a few studies have demonstrated that the use of nanostructured electrodes can improve the sensitivity of aptamer-based electrochemical sensors.^17, 19, 28^ However, the mechanism behind this enhanced sensitivity remains unclear. Investigators have attributed the enhancement to simple geometric effects due to increased surface area from the nanostructures, but without a complete picture of the underlying mechanism, optimization of the design and manufacture of such aptamer-based sensors remains challenging.

In this work, we demonstrate that an electrochemical aptamer sensor with nanoporous electrodes can achieve an up to 24-fold boost in signal and nearly four-fold improved LOD relative to an equivalent sensor employing planar electrodes. We subsequently propose and experimentally validate a mechanism underlying this improved signal output and sensitivity. In our model, these improvements result from weakened charge screening within the nanoporous electrode structure, enabling more efficient electron transfer between the redox-tagged aptamer and the gold electrode. Based on this mechanism and our testing of different nanoporous electrode structures, we demonstrate the capability to tune the electrochemical sensors in terms of signal gain, LOD, or other performance metrics. The mechanistic principles identified in this work should be broadly applicable for improving the sensitivity of aptamer-based electrochemical biosensors.

## RESULTS AND DISCUSSION

### Characterization of aptamer-immobilized nanoporous electrodes

As a proof-of-concept experiment for elucidating mechanisms of aptamer-electrode interactions within nanostructured electrodes, we employed a sensor system in which we immobilized a well-characterized doxorubicin (DOX) aptamer^29-31^ onto both planar and nanoporous gold electrodes via gold-thiol interaction (**Figure 1A**). In order to generate an electrochemical readout, the DOX aptamer was coupled to a methylene blue (MB) redox reporter. In the absence of target, the aptamer is generally in an unfolded state, limiting electron transfer between the reporter and the electrode. Target binding causes the aptamer to adopt a folded conformation, which brings the MB tag into closer proximity to the electrode surface and thus increases the electron transfer rate.

**Figure 1.**
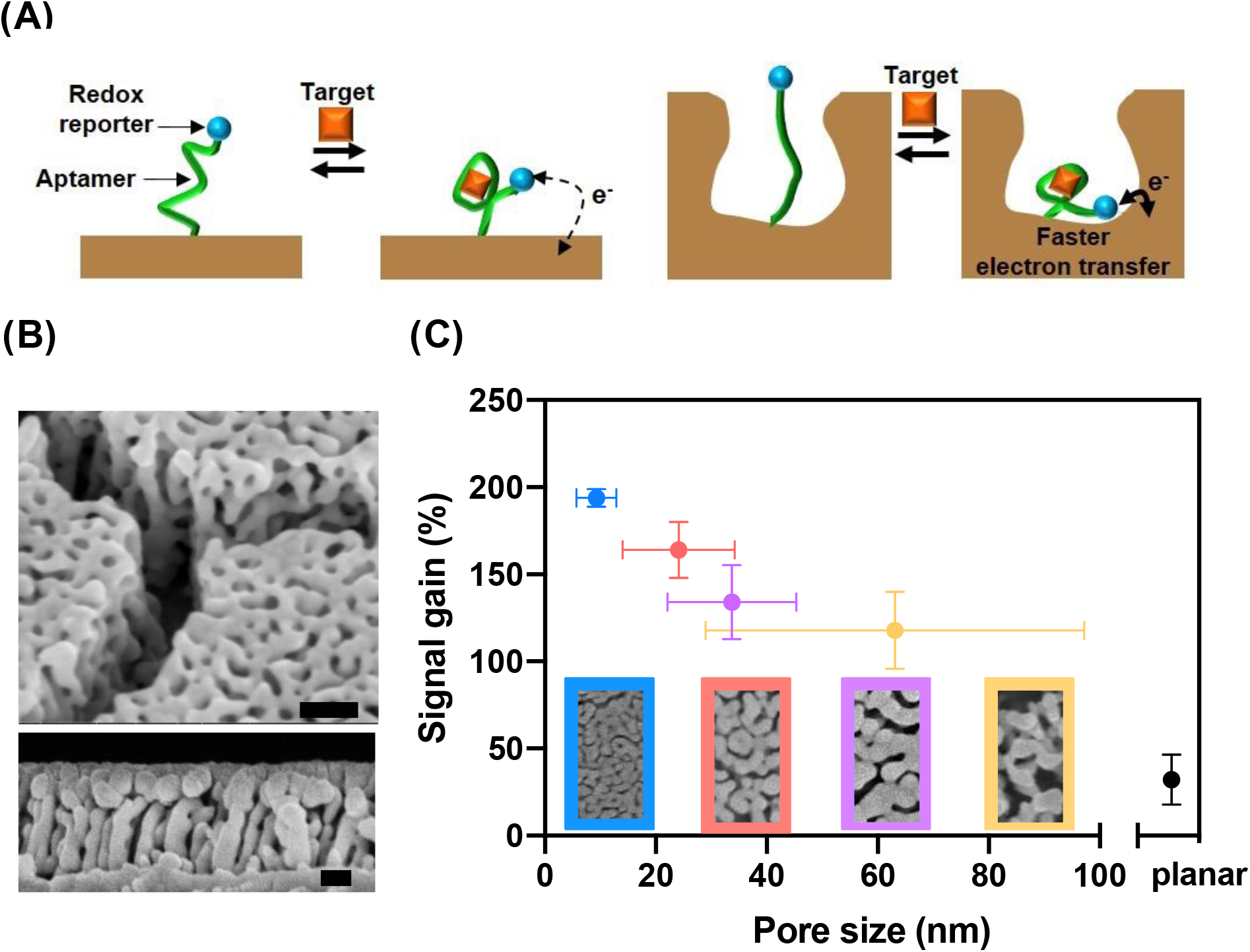
(A) Schematics of electrochemical aptamer sensors on planar (left) and nanoporous (right) electrodes. The structure-switching aptamer, end-labeled with a methylene blue (MB) reporter, is unfolded in the absence of its target, doxorubicin (DOX). This situates MB far from the electrode, yielding minimal signal. DOX binding induces aptamer folding, bringing MB close to the electrode and producing an increase in current. (B) Scanning electron microscopy (SEM) images of the nanoporous electrode. Upper and lower panels show top and side views, with scale bars of 100 nm and 50 nm, respectively. (C) Impact of pore size on signal gain. Bottom panels show SEM images of the various nanoporous electrodes. Error bars in the x-direction represent the standard deviation of different nanopores captured in the corresponding SEM images. Error bars in the y-direction represent the standard deviation of the signal gain of three replicates.

We measured the signal from each electrode using square-wave voltammetry (SWV) as an indicator of conformational change of the aptamer upon binding DOX. SWV is widely used for electrochemical aptamer sensors, as it provides higher sensitivity and lower background than most other electrochemical techniques.^32, 33^ SWV works by applying a series of small voltage steps to create electric fields in the electrochemical cell. A single square voltage waveform is applied to create two phases at each step; the positive phase partially oxidizes the MB group coupled to the aptamer, and the negative phase reduces it. The current is measured near the end of each phase, and these currents are then subtracted from each other. Because the oxidative and reductive currents have opposite signs, this subtraction maximizes the faradaic current considered in the measurement. This difference between the two phases is greatest near MB’s redox potential (E^0^ = -0.21 V *vs*. Ag/AgCl), creating a distinctive peak (**Figure S1**). We characterize these peaks using two key metrics. The signal level reflects the absolute height of the SWV peak at a defined concentration, whereas the signal gain is the ratio between the signal level at a defined concentration and a target-free control. Higher signal level helps distinguish SWV peaks from noise, while higher signal gain makes it easier to quantify different concentrations.

We fabricated the nanoporous electrodes using a dealloying process with Ag:Au alloys.^34, 35^ The pore size in these electrodes can be tuned by adjusting the ratio of Ag to Au. We used two approaches—thermal annealing^26^ and electrochemical coarsening^27^—to adjust the average pore and ligament sizes in our nanoporous structures. Using these two mechanisms, we were able to fabricate nanoporous electrodes with average pore sizes between 9.3 nm and 63.1 nm (**Figure 1B**). For comparison, we also fabricated planar electrodes with the same footprint (100 μm by 100 μm). By controlling the nanopore size, we found that we could engineer the signal gain and improve the signal level of our sensor. We characterized the signal gain of nanoporous electrodes with different pore sizes after adding a saturating concentration (100 µM) of DOX. Decreasing the pore size led to higher signal gain, where the smallest nanopores showed the highest signal gain of 194% versus 32% for the planar electrode (**Figure 1C**). We optimized for signal gain by utilizing the smallest average nanopore size (9.3 nm) for subsequent experiments.

Next, we optimized several of the control parameters for our nanoporous sensor. Changing the concentration of aptamer molecules applied during the immobilization process offers a way to control the probe density and inter-aptamer spacing on the electrode surface.^36, 37^ High aptamer density leads to steric hindrance, which impedes target binding, whereas excessively low aptamer density can cause the signal level to decrease to undetectable levels. We prepared nanoporous and planar electrodes using five different aptamer concentrations (0.1 μM, 0.5 μM, 1 μM, 3 μM, 10 μM), yielding molecular probe densities that we labeled as d0.1, d0.5, d1, d3, and d10, respectively (refer to **Figure S3** for details). Then we generated calibration curves by measuring SWV at several concentrations of DOX, ranging from 100 nM to 100 μM. In parallel, we also optimized the frequency and amplitude of the waveform applied to the electrochemical cell during measurement, parameters that affect transduction from the redox reporter to the electrode.^38^ For each aptamer concentration, we performed SWV with five different frequencies (50, 100, 200, 300, and 400 Hz) and three different amplitudes (20, 50, and 100 mV) (refer to **Figure S4** for details).

The nanoporous electrodes consistently produced greatly improved performance relative to planar electrodes. We optimized both the nanoporous and planar sensor for the abovementioned conditions across several metrics: signal gain, signal level, and LOD (**Table 1)**. At 10 μM DOX— the upper limit of the clinically-relevant range for this drug^30^—we could independently improve signal gain by as much as three-fold (from 59% to 179%) and signal level by as much as 24-fold (from 30.4 nA to 728.0 nA) for the nanoporous electrodes relative to their planar counterparts. This superior performance can be partly explained by the much larger surface area achieved with the nanoporous electrode while keeping the same footprint as a planar electrode. This allows more total probes to be immobilized, which generates higher currents. At the same time, having more probes on the electrode includes more binding events in the ensemble measurement, lowering the variance of the measured signal. This is reflected in the smaller error between nanoporous replicates (∼4% CV) versus planar replicates (∼20% CV) (**Figure 2A**; refer to **Table S2** for details). In combination with the increased signal gain that we observed for our electrodes, the lower variation and increased signal level result in a decreased LOD (**Figure 2B**). Indeed, by leveraging the additional signal gain and signal level, the sensor LOD could be decreased nearly four-fold, from 101.3 ± 16.8 nM on planar electrodes to 28.5 ± 1.4 nM on nanoporous electrodes.

**Table 1.** Conditions yielding maximum sensor signal gain, signal level, and LOD for nanoporous and planar electrodes. Signal gain and level measurements are for 10 μM DOX; enhancement ratio describes improved performance in each optimized metric for nanoporous versus planar sensors.

| Optimized metrics | Electrode | Probe density | SWV Hz | SWV Amp | Signal gain | Signal level (nA) | LOD ($\mu\text{M}$) | Enhancement ratio |
| --- | --- | --- | --- | --- | --- | --- | --- | --- |
| Signal gain | nanoporous | d1 | 400hz | 20mV | 179.00% | 180.9 | 0.080 | 3.0 |
|  | planar | d1 | 200hz | 20mV | 59.37% | 2.7224 | 0.273 |  |
| Signal level | nanoporous | d10 | 400hz | 100mV | 76.14% | 727.99 | 0.116 | 23.9 |
|  | planar | d10 | 400hz | 100mV | 15.26% | 30.415 | 0.107 |  |
| LOD | nanoporous | d1 | 300hz | 20mV | 157.69% | 143.72 | 0.028 | 3.6 |
|  | planar | d3 | 100hz | 50mV | 30.78% | 3.1551 | 0.100 |  |

**Figure 2.**
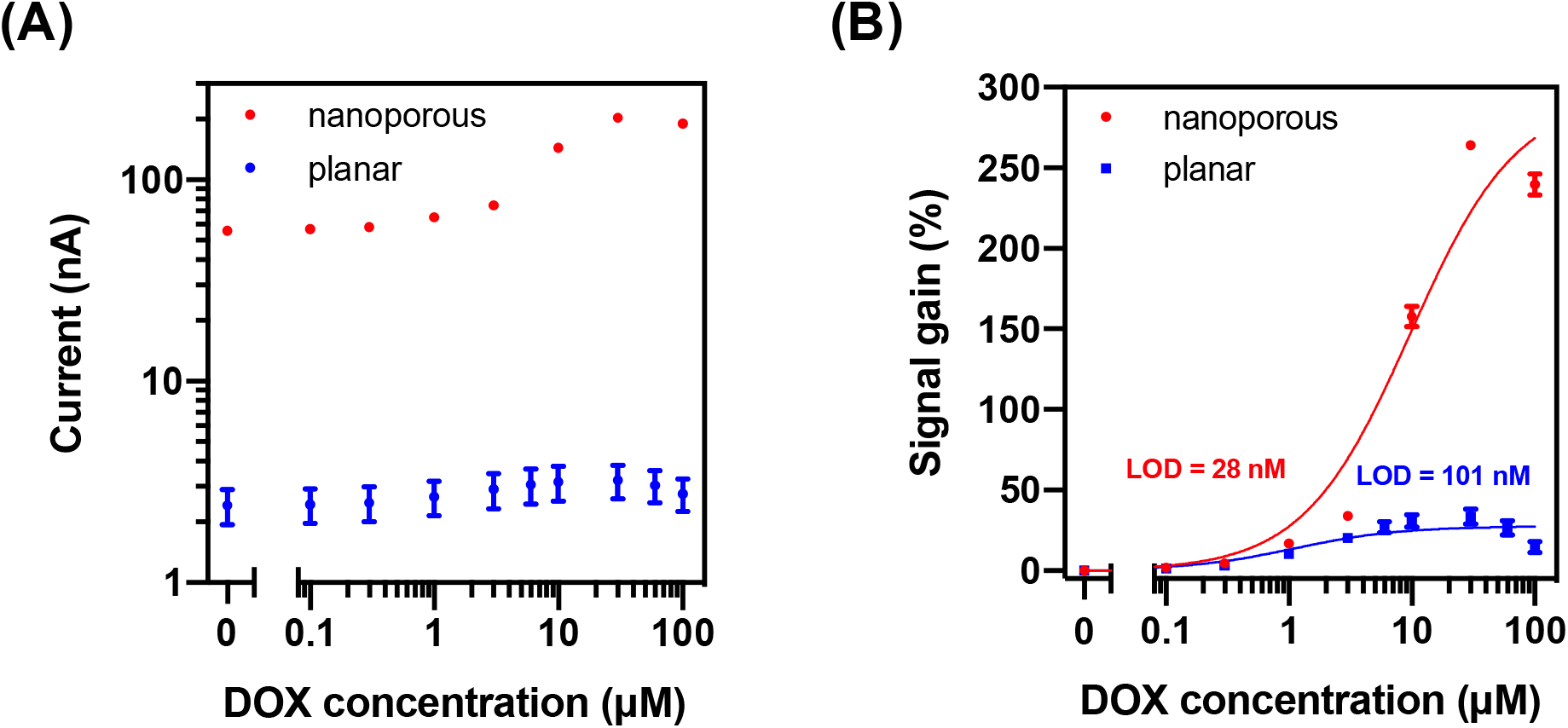
(A) Signal level and (B) signal gain of electrochemical aptamer sensors employing nanoporous (red) and planar (blue) electrodes in the presence of 100 nM–100 μM DOX. Plots are averaged over three replicates. Error bars represent the standard deviation.

### Mechanistic explanation of signal enhancement

A number of researchers have reported increases in signal gain with nanostructured electrodes^17, 19, 28^, but without a definitive mechanistic explanation for this phenomenon. Some studies have suggested that the increased sensitivity is due to the change in the “effective K_D_” of the molecular probe, resulting from local increases in analyte concentration^15, 26^. Although this maybe valid under certain conditions^39^, our data from nanoporous and planar electrodes showed increased signal gain without a significant change in binding thermodynamics (K_D_ = 1.71 μM and 2.14 μM, respectively).

As an alternative, we hypothesized that the nanoporous structure of our electrodes was directly affecting the kinetics of electron transfer between the MB reporter and the electrode through its weakening effect on charge screening. In electrochemical aptamer sensors that use SWV, electric fields are applied across the electrochemical cell to initiate concentration-dependent electron transfer.^40, 41^ However, in physiological samples and other electrolytic solutions, these electric fields are confined to the EDL, a small region adjacent to the electrode where ions screen the field being applied. This EDL is quite small—on the order of the Debye length (<1 nm)—and defines a small volume that the MB reporter must enter for electron transfer to occur. This can be approximated in terms of the Debye volume: the space between the electrode surface and an imaginary surface one Debye length normal to it. It has been shown that limiting the Debye volume at an electrode surface extends the EDL farther into the solution, thereby lessening the extent of electric field screening at that interface.^42, 43^ Consequently, the stronger electric fields within the nanopores increase the probability of a faradaic electron transfer event for a given conformation of an aptamer probe. Indeed, because of their high density of nanoscale features, nanoporous electrodes offer exactly the type of interface where screening is weaker and where we would predict faradaic electron transfer to be accelerated (**Figure 3**).

**Figure 3.**
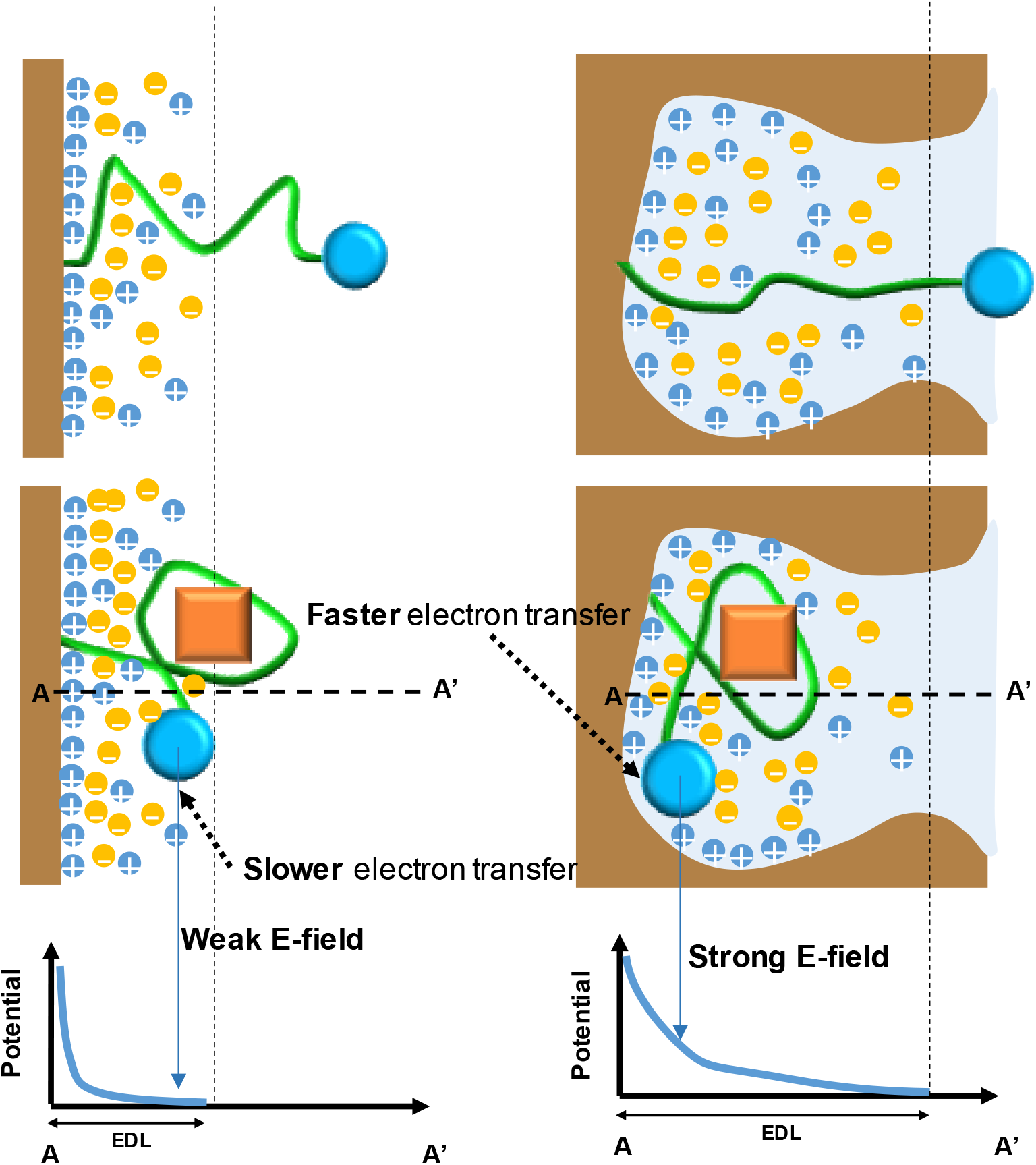
Scheme of how the electric double layer (EDL) from a planar (left) and nanoporous electrode (right) affects electron transfer from a MB reporter (blue circle) tethered to the aptamer (green) before and after target (orange square) binding. During electrochemical measurement, the MB reporter interacts with the EDL (shaded blue region), where a closer distance between the reporter and the electrode surface leads to faster electron transfer. In the nanoporous electrode, the MB group experiences stronger electric fields. A and A’ represent the electrode surface and the maximum distance of MB from the electrode, respectively.

We carried out a 2D simulation study in COMSOL to test this hypothesis. Because the morphology of nanoporous electrodes is irregular and highly variable, we simplified our study by simulating several basic geometries that we believe are representative of geometric elements typically found in true nanoporous electrode structures. We focused on semicircles with radii of 5, 20, and 50 nm (**Figure 4A**) and triangles with base widths and heights of 5, 20, and 50 nm (**Figure 4B–C**). We applied 100 mV to the interfaces under study, approximating the screening conditions in the cell at the redox potential of MB. For each structure, the electric potential was extracted for a 10-nm straight line from the structure’s apex to evaluate potential decay in space. Typically, any applied potential will decay exponentially with distance, with a spatial rate constant defined by the Debye length. However, our simulation suggested that nanoporous gold surfaces could diminish the effects of screening, depending on the size of the cavities involved. As critical dimensions decreased in size, the potential decayed less sharply with distance, indicating that the EDL is extended in the nanostructure and that screening is weakened. Semicircles exhibited stronger screening than triangular shapes, and this is likely due to the sides of the triangles creating closer distances between adjacent surfaces, mimicking nano-gap structures.^44^ Indeed, our simulation results provide strong evidence that the spatial scale of the nanopore structures in our electrodes is sufficient to support weakened charge screening relative to a planar surface.

**Figure 4.**
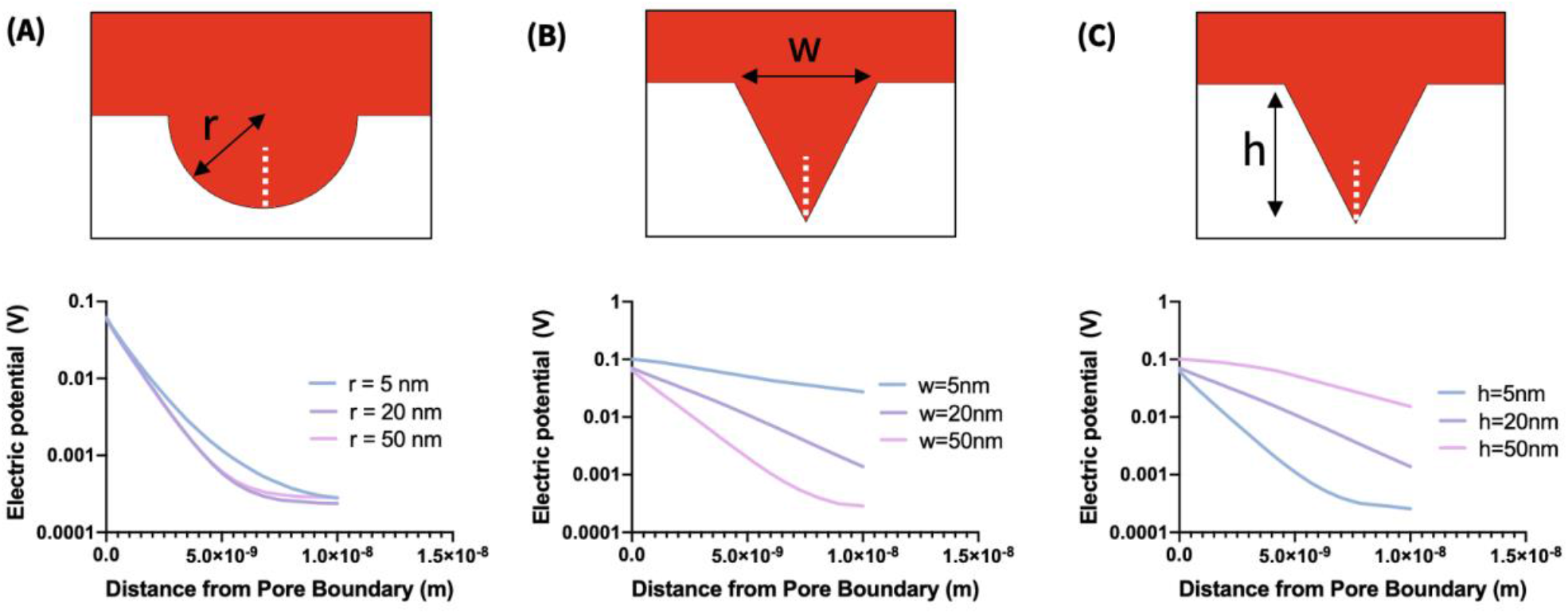
Simulations of screening for nanostructured electrode geometries with 100 mV applied. Top panels show the tested geometries, including (A) semicircular pores of varying radius, and triangular pores of (B) fixed height (20 nm) and varying width (w) or (C) fixed width (20 nm) and varying height (h). Bottom panels show electric potentials for a 10-nm cut line extending vertically from the deepest point in the structure (white dashed line in top panels).

To experimentally test the effect of weakened screening on electrochemical measurements, we varied the ionic strength of the sample and generated calibration curves by performing SWV on the aptamer-functionalized nanoporous electrodes. Ionic strength is known to act as a ‘control knob’ for screening at electrode-electrolyte interfaces, where the relationship between the ionic strength (*I*) and Debye length (*λ*_*D*_) is 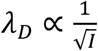. Thus, by tuning the ionic strength of the sample, we can modulate the screening conditions at the electrode interface to study how screening affects electron transfer from MB to the electrode (**Figure 5A)**. We prepared samples containing various concentrations of DOX in 0.1X, 1X, or 10X SSC buffer, where 1X SSC buffer contains 150 mM NaCl and 15 mM trisodium citrate (pH 7.0). We then measured square-wave voltammograms with planar (**Figure 5B**) and nanoporous electrodes (**Figure 5C**) in each sample to generate a calibration curve for each buffer concentration (refer to **Figure S5** for details of the effect of ionic strength on measurement). We note that at high analyte concentrations, we observed a decreasing signal gain under some experimental conditions. A similar trend was also observed in a recent study, and this is likely due to the interference of the target with electron transfer between the target-aptamer complex and the electrode surface.^28^

**Figure 5.**
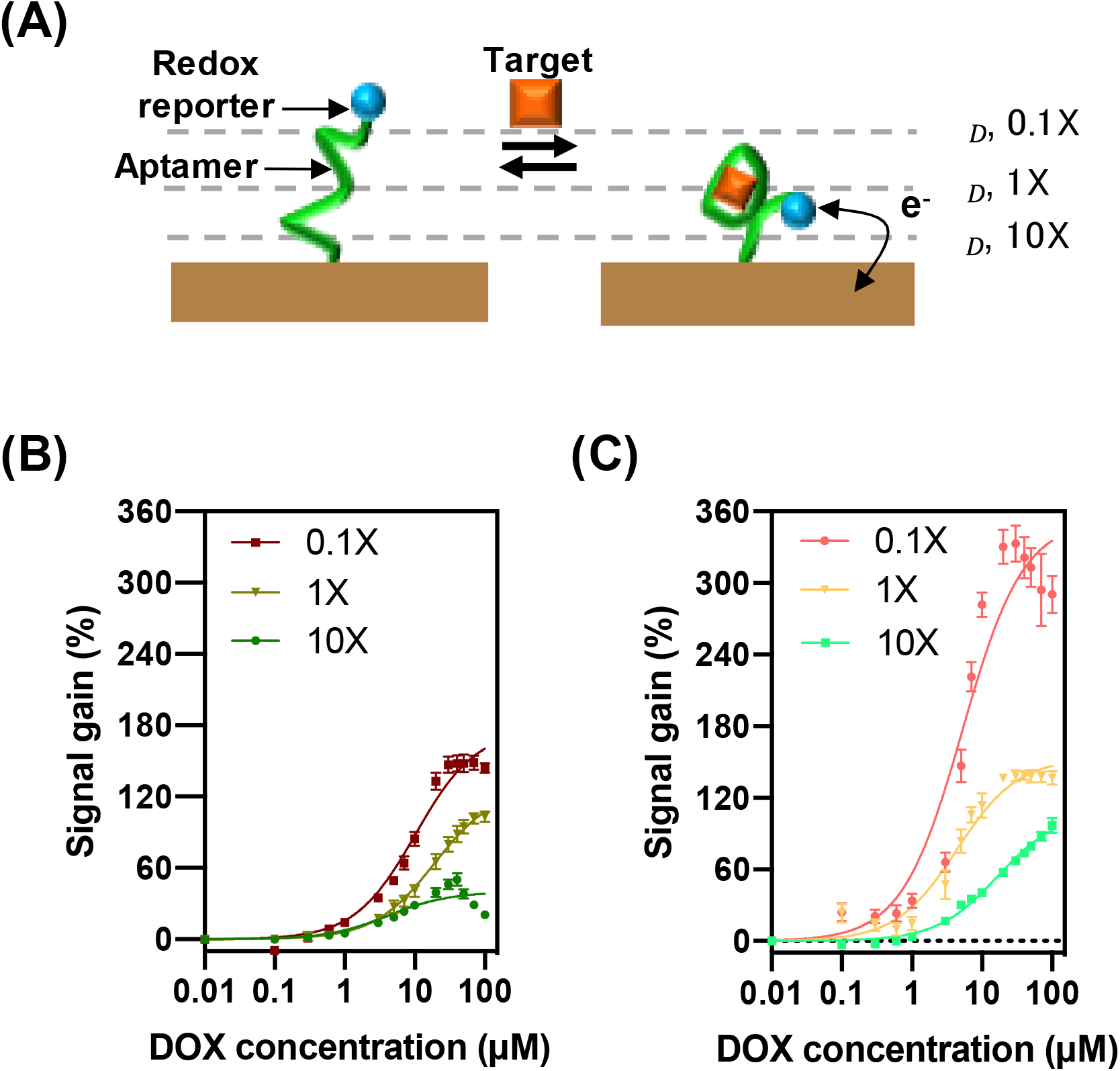
(A) Ionic strength affects the EDL configuration, which affects transduction between the MB reporter and the electrode. As the ionic strength of the solution increases, the Debye length (*λ*_*D*_) decreases. Calibration curves were generated by performing SWV on aptamer-functionalized (B) planar and (C) nanoporous electrodes with samples spiked with varying concentrations of DOX in different dilutions of SSC buffer to assess the impact of ionic strength on signal gain. Datapoints are averaged over three replicates, and error bars show standard deviation.

These calibration curves confirmed our theory—as the ionic strength of the solution decreases, the EDL extends farther into the solution and signal gain increases for both planar and nanoporous electrodes. This is consistent with our simulations, which showed that the EDL extends farther in smaller geometries (**Figure 4**), and our data showing that nanoporous electrodes with smaller pores generate higher signal gain (**Figure 1C**). Notably, the effective *K*_*D*_ of the sensor was also affected by changes in ionic strength. This effect is likely a consequence of charge screening as well, because electrostatic changes alter the intramolecular (*e*.*g*., energetics of Watson-Crick base pairing, formation of secondary and tertiary structures) and intermolecular interactions (*e*.*g*., binding energy) that determine the thermodynamics of the aptamer sensor.

Finally, we further validated our hypothesis by generating chronoamperograms to show that weakened screening in the nanoporous electrodes indeed causes accelerated electron transfer. We tested aptamer-functionalized nanoporous electrodes with several concentrations of DOX up to 100 μM, averaging 50 measurements from each electrode to reduce noise at the lower current ranges. These chronoamperogram measurements are made up of two types of current: non-faradaic current, which captures the movement of ions to charge the EDL, and faradaic current, which captures the electrochemical transfer of charge to or from redox reporters. The total current can be modeled as a two-phase exponential decay, where the faster phase (*τ*_*fast*_) represents the capacitive EDL charging current and the slower phase (*τ*_*slow*_) represents the electrochemical current (**Figure 6A**).^45^ With increasing concentrations of DOX, we expect more aptamers to be bound to their target, and we predictably observed that *τ*_*slow*_ became shorter with increasing target concentrations. This indicates that more charge is transferred sooner as the aptamer binds to its target. We found that *τ*_*slow*_ was consistently shorter across DOX concentrations for nanoporous electrodes compared to planar electrodes, indicating faster faradaic reactions between the redox reporter and electrode (**Figure 6B**).

**Figure 6.**
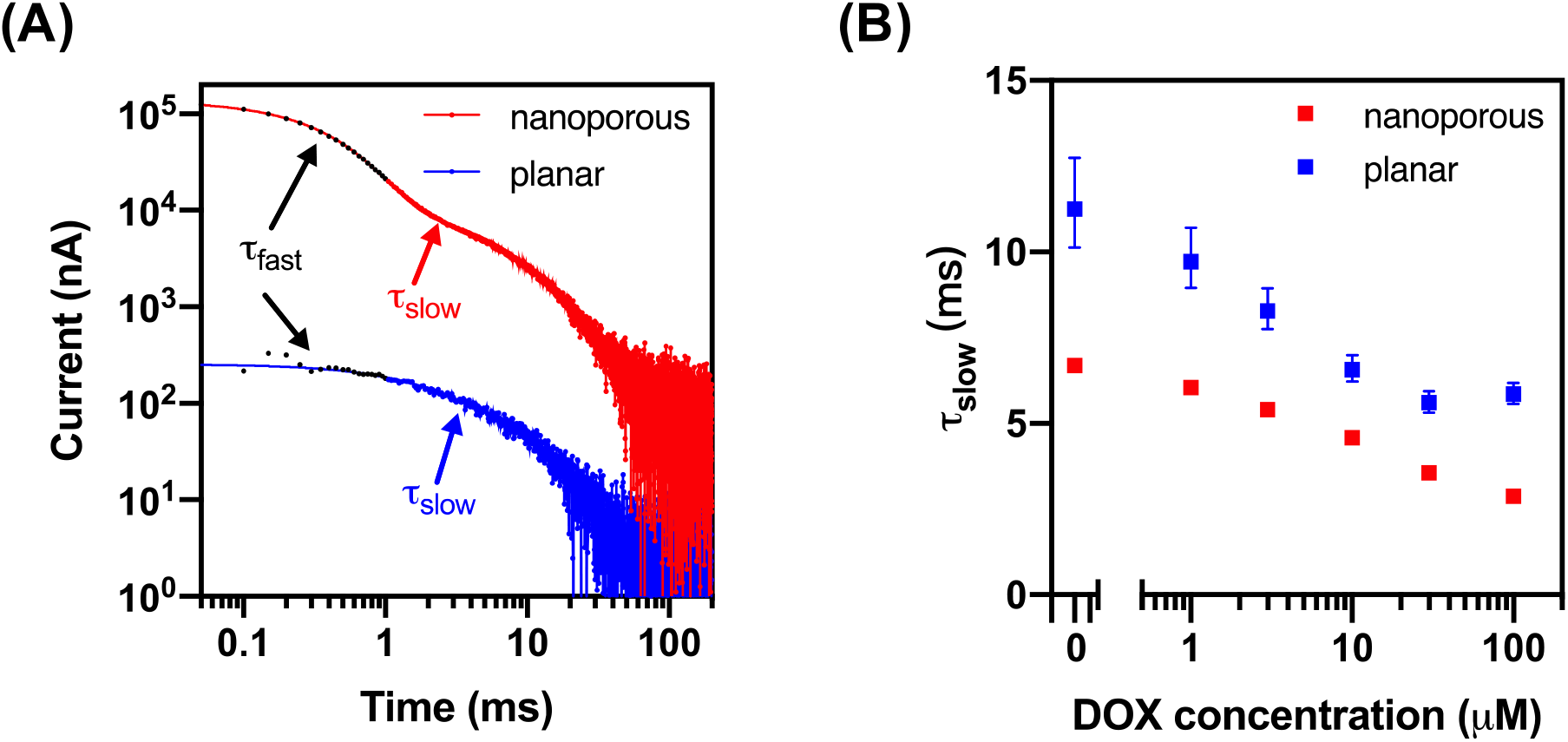
(A) Chronoamperograms of planar (blue) and nanoporous (red) electrode sensors with an electrode size of 500 μm by 500 μm in the absence of DOX (log scale, negative values not shown). (B) The decay time (*τ*_*slow*_) of both electrodes at various DOX concentrations. Error bars represent the confidence intervals of the curve fitting.

This observation is consistent with the core mechanism of structure-switching electrochemical aptamer sensors, in which charges are exchanged when the redox reporter encounters sufficiently high electric fields within the electrode’s EDL. If a binding event brings the MB reporter closer to the surface (on average), the faradaic reaction will complete faster. Importantly, our data demonstrates that the extended EDL on the nanoporous electrodes also facilitates more efficient electron transfer than on the planar electrodes.

## CONCLUSION

In this work, we have developed and experimentally validated a model to explain why nanoporous gold electrodes can deliver considerably improved performance relative to planar gold surfaces in the context of electrochemical aptamer sensors. We first showed that nanoporous electrodes consistently offer superior signal output and signal gain relative to planar surfaces in an electrochemical aptamer sensor for the chemotherapeutic drug doxorubicin, and show that these electrodes can be optimized to achieve ∼24-fold higher signal level and approximately 4-fold lower LOD relative to planar gold electrodes. We subsequently hypothesized that this greater sensitivity and signal output may be the consequence of the decreased Debye volume and reduced impact of charge screening within these nanopores, and were able to test and confirm this hypothesis via both computer simulations and experimental testing. Collectively, our results reveal design principles that can guide the production of electrochemical aptamer sensors with optimized detection performance—for example, employing smaller nanopores to maximize the signal gain generated in response to target binding, or tuning the square-wave voltammetry settings to achieve the best LOD. We believe that this ability to engineer electron transfer efficiency should prove highly valuable for improving the performance of electrochemical biosensors for a wide range of biomolecules in diagnostic and health monitoring applications.

## METHODS

### Reagents and materials

The DOX aptamer was adapted from our previous studies^30, 31^ and obtained from Biosearch Technologies: 5’-HS-C6-ACCATCTGTGTAAGGGGTAAGGGGTGGT-MB-3’, where MB indicates the methylene blue reporter. 6-mercapto-1-hexanol (6-MCH), tris(2-carboxyethyl)phosphine (TCEP), and doxorubicin (DOX) were obtained from Sigma-Aldrich. A 1X stock solution of saline-sodium citrate (SSC) buffer was prepared by dilution of SSC stock solution (20X, pH 7.4, Thermo Fisher Scientific) with nuclease-free water. DOX solution was prepared before measurement in SSC buffer.

### Device fabrication and characterization

A 100 μm x 100 μm Ti/Au/Ag:Au (10 nm/50 nm/300 nm) film was patterned onto glass slides via lift-off process. The ratio of co-sputtered Ag:Au film was 2:1.^26, 27^ Next, Ag was selectively etched by nitric acid (70% v/v) for 5 minutes, forming the Au nanoporous microelectrode. Finally, the whole area except the 100 μm x 100 μm Au nanoporous microelectrode was encapsulated.

Post-treatment of the fabricated nanoporous electrode slide included either a thermal annealing process or an electrochemical coarsening process (**Table S1**). The former process entails treating the nanoporous electrode slide at 230 °C for 10 min on a hotplate. This resulted in larger nanopores and fewer cracks; the average pore size was 9.29 nm at room temperature (RT), but increased to 24.12 nm at 230 °C. Electrochemical coarsening is achieved through cyclic voltammetry (CV) in 0.5 M sulfuric acid solution with a potential window of 0.3 V to 1.2 V for multiple scans. The resulting nanostructures were characterized by scanning electron microscopy (FEI Nova NanoSEM 450). The surface of the nanoporous gold was repetitively oxidized and reduced to form larger pore sizes ranging from 9.29 nm to 33.7 nm. By performing both thermal annealing and electrochemical coarsening, we could increase the pore size up to 63.11 nm. CV of nanoporous and planar electrodes was conducted in 0.05 M sulfuric acid at a scan rate of 50 mV/s over the potential range −0.25 to +1.75 mV to determine the effective surface area of the electrode (**Figure S2**).^26^

### Electrochemical aptamer sensor characterization

After rinsing the nanoporous electrode slide with deionized (DI) water, the freshly prepared slide was incubated with 1 μM aptamer in 1X SSC buffer for one hour. Soft-polymer PDMS wells were sealed on top of the nanoporous electrode slide to hold each electrode’s solution. To study the effects of aptamer surface density, we also prepared other concentrations of DOX aptamer. Before incubation, 100 μM aptamer in DI water was reacted with a 1,000-fold excess of TCEP solution to produce free thiol groups for aptamer immobilization. After immobilization, the nanoporous electrode slide was washed with excess buffer and then incubated with 10 mM 6-MCH for two hours to passivate the remaining electrode surface. The electrode was then stored in 1X SSC before electrochemical measurement.

All measurements were performed in PDMS wells with a PalmSens 4 potentiostat. All working electrodes—whether nanoporous or planar—were 100 μm by 100 μm unless specified otherwise. Commercial Ag/AgCl reference electrodes and Pt wire counter electrodes were from CH Instruments. SWV was carried out in 1× SSC buffer over the potential range of -0.5 V to 0.0 V with an amplitude of 20–100 mV, step-size of 2 mV, and pulse frequencies ranging from 50–400 Hz. DOX binding curves were generated across various probe densities and SWV parameters as part of our optimization. We also calculated the LOD for DOX based on the concentration that gave a signal three baseline standard deviations (σ_b_) above zero:

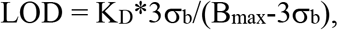

where K_D_ and B_max_ are the dissociation constant and maximum specific binding, respectively. These values were obtained by fitting the signal gain at different DOX concentrations to a Langmuir isotherm.^46^

### COMSOL simulation methodology

Simulations were done in COMSOL 5.5 using the Electrochemistry module. By coupling the “electrostatics” and “transport of dilute species” physics, the Poisson-Nernst-Planck system of equations was solved for a 100 mM NaCl electrolyte with Debye length ∼1 nm. Boundary conditions were no flux at the interface under study, to define the electrode as polarizable and eliminate effects of any faradaic reactions, and equilibrium concentrations were defined to mimic a controlled potential state as established by a potentiostat or similar circuitry. Shapes were simulated as part of a large square field with a side length of 1 micrometer, to prevent any effects due to other boundaries.

## Supporting information

Fu_Seo_Kesler_SI

## ACKNOWLEDGEMENTS

This work was supported by the Chan-Zuckerberg Biohub, the Biomedical Advanced Research and Development Agency (BARDA, 75A50119C00051) and the National Institutes of Health (NIH, OT2OD025342). We thank Ian Thompson and Jun-Chau Chien for their thoughtful comments and valuable suggestions on the experiments.

## AUTHOR CONTRIBUTION

K.F., J.W.S. and H.T.S. initiated the project. K.F. and J.W.S. designed and characterized the device. K.F., J.W.S. and V.K. designed the experiments. K.F. executed the experiments and collected the data. V.K. conceived the theoretical model and performed the simulations. K.F. and V.K. analyzed the data. K.F., V.K., J.W.S. and H.T.S. wrote the paper. All authors discussed the data and edited the paper.

## Notes

### Competing Interest Statement

The authors have declared no competing interest.

## REFERENCES

1. Rothe, J., Frey, O., Stettler, A., Chen, Y., Hierlemann, A., Fully integrated CMOS microsystem for electrochemical measurements on 32 × 32 working electrodes at 90 frames per second. Analytical Chemistry 2014, 86 (13), 6425–6432.

2. Bellin, D. L., Sakhtah, H., Rosenstein, J. K., Levine, P. M., Thimot, J., Emmett, K., Dietrich, L. E. P., Shepard, K. L., Integrated circuit-based electrochemical sensor for spatially resolved detection of redox-active metabolites in biofilms. Nature Communications 2014, 5 (1), 3256.

3. Slinker, J. D., Muren, N. B., Gorodetsky, A. A., Barton, J. K., Multiplexed DNA-Modified Electrodes. Journal of the American Chemical Society 2010, 132 (8), 2769–2774.

4. Pan, L., Yu, G., Zhai, D., Lee, H. R., Zhao, W., Liu, N., Wang, H., Tee, B. C. K., Shi, Y., Cui, Y., Bao, Z., Hierarchical nanostructured conducting polymer hydrogel with high electrochemical activity. Proceedings of the National Academy of Sciences 2012, 109 (24), 9287.

5. Gao, W., Emaminejad, S., Nyein, H. Y. Y., Challa, S., Chen, K., Peck, A., Fahad, H. M., Ota, H., Shiraki, H., Kiriya, D., Lien, D.-H., Brooks, G. A., Davis, R. W., Javey, A., Fully integrated wearable sensor arrays for multiplexed in situ perspiration analysis. Nature 2016, 529 (7587), 509–514.

6. Wang, S., Zhang, L., Wan, S., Cansiz, S., Cui, C., Liu, Y., Cai, R., Hong, C., Teng, T., Shi, M., Wu, Y., Dong, Y., Tan, W., Aptasensor with Expanded Nucleotide Using DNA Nanotetrahedra for Electrochemical Detection of Cancerous Exosomes. ACS Nano 2017, 11 (4), 3943–3949.

7. Bandodkar, A. J., Gutruf, P., Choi, J., Lee, K., Sekine, Y., Reeder, J. T., Jeang, W. J., Aranyosi, A. J., Lee, S. P., Model, J. B., Ghaffari, R., Su, C.-J., Leshock, J. P., Ray, T., Verrillo, A., Thomas, K., Krishnamurthi, V., Han, S., Kim, J., Krishnan, S., Hang, T., Rogers, J. A., Battery-free, skin-interfaced microfluidic/electronic systems for simultaneous electrochemical, colorimetric, and volumetric analysis of sweat. Science Advances 2019, 5 (1), eaav3294.

8. Kim, J., Campbell, A. S., de Ávila, B. E.-F., Wang, J., Wearable biosensors for healthcare monitoring. Nature Biotechnology 2019, 37 (4), 389–406.

9. Patolsky, F., Lichtenstein, A., Willner, I., Detection of single-base DNA mutations by enzyme-amplified electronic transduction. Nature Biotechnology 2001, 19 (3), 253–257.

10. Rissin, D. M., Kan, C. W., Campbell, T. G., Howes, S. C., Fournier, D. R., Song, L., Piech, T., Patel, P. P., Chang, L., Rivnak, A. J., Ferrell, E. P., Randall, J. D., Provuncher, G. K., Walt, D. R., Duffy, D. C., Single-molecule enzyme-linked immunosorbent assay detects serum proteins at subfemtomolar concentrations. Nature Biotechnology 2010, 28 (6), 595–599.

11. Hu, J., Wang, T., Kim, J., Shannon, C., Easley, C. J., Quantitation of Femtomolar Protein Levels via Direct Readout with the Electrochemical Proximity Assay. Journal of the American Chemical Society 2012, 134 (16), 7066–7072.

12. Labib, M., Khan, N., Ghobadloo, S. M., Cheng, J., Pezacki, J. P., Berezovski, M. V., Three-Mode Electrochemical Sensing of Ultralow MicroRNA Levels. Journal of the American Chemical Society 2013, 135 (8), 3027–3038.

13. Kelley, S. O., Mirkin, C. A., Walt, D. R., Ismagilov, R. F., Toner, M., Sargent, E. H., Advancing the speed, sensitivity and accuracy of biomolecular detection using multi-length-scale engineering. Nature Nanotechnology 2014, 9 (12), 969–980.

14. Tavallaie, R., McCarroll, J., Le Grand, M., Ariotti, N., Schuhmann, W., Bakker, E., Tilley, R. D., Hibbert, D. B., Kavallaris, M., Gooding, J. J., Nucleic acid hybridization on an electrically reconfigurable network of gold-coated magnetic nanoparticles enables microRNA detection in blood. Nature Nanotechnology 2018, 13 (11), 1066–1071.

15. Soleymani, L., Fang, Z., Sargent, E. H., Kelley, S. O., Programming the detection limits of biosensors through controlled nanostructuring. Nature Nanotechnology 2009, 4, 844.

16. Huang, X.-J., O’Mahony, A. M., Compton, R. G., Microelectrode Arrays for Electrochemistry: Approaches to Fabrication. Small 2009, 5 (7), 776–788.

17. Liu, J., Wagan, S., Dávila Morris, M., Taylor, J., White, R. J., Achieving Reproducible Performance of Electrochemical, Folding Aptamer-Based Sensors on Microelectrodes: Challenges and Prospects. Analytical Chemistry 2014, 86 (22), 11417–11424.

18. Wang, T., Viennois, E., Merlin, D., Wang, G., Microelectrode miRNA Sensors Enabled by Enzymeless Electrochemical Signal Amplification. Analytical Chemistry 2015, 87 (16), 8173–8180.

19. Arroyo-Curras, N., Scida, K., Ploense, K. L., Kippin, T. E., Plaxco, K. W., High Surface Area Electrodes Generated via Electrochemical Roughening Improve the Signaling of Electrochemical Aptamer-Based Biosensors. Analytical Chemistry 2017, 89 (22), 12185–12191.

20. Hu, K., Lan, D., Li, X., Zhang, S., Electrochemical DNA Biosensor Based on Nanoporous Gold Electrode and Multifunctional Encoded DNA−Au Bio Bar Codes. Analytical Chemistry 2008, 80 (23), 9124–9130.

21. Oja, S. M., Wood, M., Zhang, B., Nanoscale Electrochemistry. Analytical Chemistry 2013, 85 (2), 473–486.

22. Sage, A. T., Besant, J. D., Lam, B., Sargent, E. H., Kelley, S. O., Ultrasensitive Electrochemical Biomolecular Detection Using Nanostructured Microelectrodes. Accounts of Chemical Research 2014, 47 (8), 2417–2425.

23. Salamifar, S. E., Lai, R. Y., Fabrication of Electrochemical DNA Sensors on Gold-Modified Recessed Platinum Nanoelectrodes. Analytical Chemistry 2014, 86 (6), 2849–2852.

24. Su, S., Wu, Y., Zhu, D., Chao, J., Liu, X., Wan, Y., Su, Y., Zuo, X., Fan, C., Wang, L., On-Electrode Synthesis of Shape-Controlled Hierarchical Flower-Like Gold Nanostructures for Efficient Interfacial DNA Assembly and Sensitive Electrochemical Sensing of MicroRNA. Small 2016, 12 (28), 3794–3801.

25. Fu, K., Kwon, S.-R., Han, D., Bohn, P. W., Single Entity Electrochemistry in Nanopore Electrode Arrays: Ion Transport Meets Electron Transfer in Confined Geometries. Accounts of Chemical Research 2020, 53 (4), 719–728.

26. Daggumati, P., Matharu, Z., Seker, E., Effect of Nanoporous Gold Thin Film Morphology on Electrochemical DNA Sensing. Analytical Chemistry 2015, 87 (16), 8149–8156.

27. Matharu, Z., Daggumati, P., Wang, L., Dorofeeva, T. S., Li, Z., Seker, E., Nanoporous-Gold-Based Electrode Morphology Libraries for Investigating Structure–Property Relationships in Nucleic Acid Based Electrochemical Biosensors. ACS Applied Materials & Interfaces 2017, 9 (15), 12959–12966.

28. Li, S., Lin, L., Chang, X., Si, Z., Plaxco, K. W., Khine, M., Li, H., Xia, F., A wrinkled structure of gold film greatly improves the signaling of electrochemical aptamer-based biosensors. RSC Advances 2021, 11 (2), 671–677.

29. Wochner, A., Menger, M., Orgel, D., Cech, B., Rimmele, M., Erdmann, V. A., Glökler, J., A DNA aptamer with high affinity and specificity for therapeutic anthracyclines. Analytical Biochemistry 2008, 373 (1), 34–42.

30. Ferguson, B. S., Hoggarth, D. A., Maliniak, D., Ploense, K., White, R. J., Woodward, N., Hsieh, K., Bonham, A. J., Eisenstein, M., Kippin, T. E., Plaxco, K. W., Soh, H. T., Real-Time, Aptamer-Based Tracking of Circulating Therapeutic Agents in Living Animals. Science Translational Medicine 2013, 5 (213), 213ra165.

31. Mage, P. L., Ferguson, B. S., Maliniak, D., Ploense, K. L., Kippin, T. E., Soh, H. T., Closed-loop control of circulating drug levels in live animals. Nature Biomedical Engineering 2017, 1 (5), 0070.

32. Cobb, S. J., Macpherson, J. V., Enhancing Square Wave Voltammetry Measurements via Electrochemical Analysis of the Non-Faradaic Potential Window. Analytical Chemistry 2019, 91 (12), 7935–7942.

33. Holtan, M. D., Somasundaram, S., Khuda, N., Easley, C. J., Nonfaradaic Current Suppression in DNA-Based Electrochemical Assays with a Differential Potentiostat. Analytical Chemistry 2019, 91 (24), 15833–15839.

34. Erlebacher, J., Aziz, M. J., Karma, A., Dimitrov, N., Sieradzki, K., Evolution of nanoporosity in dealloying. Nature 2001, 410 (6827), 450–453.

35. Ding, Y., Erlebacher, J., Nanoporous Metals with Controlled Multimodal Pore Size Distribution. Journal of the American Chemical Society 2003, 125 (26), 7772–7773.

36. Ricci, F., Lai, R. Y., Heeger, A. J., Plaxco, K. W., Sumner, J. J., Effect of Molecular Crowding on the Response of an Electrochemical DNA Sensor. Langmuir 2007, 23 (12), 6827–6834.

37. Mahshid, S. S., Camiré, S., Ricci, F., Vallée-Bélisle, A., A Highly Selective Electrochemical DNA-Based Sensor That Employs Steric Hindrance Effects to Detect Proteins Directly in Whole Blood. Journal of the American Chemical Society 2015, 137 (50), 15596–15599.

38. Dauphin-Ducharme, P., Plaxco, K. W., Maximizing the Signal Gain of Electrochemical-DNA Sensors. Analytical Chemistry 2016, 88 (23), 11654–11662.

39. Wilson, B. D., Hariri, A. A., Thompson, I. A. P., Eisenstein, M., Soh, H. T., Independent control of the thermodynamic and kinetic properties of aptamer switches. Nature Communications 2019, 10 (1), 5079.

40. Ramaley, L., Krause, M. S., Theory of square wave voltammetry. Analytical Chemistry1969, 41 (11), 1362–1365.

41. Xiao, Y., Lubin, A. A., Heeger, A. J., Plaxco, K. W., Label-Free Electronic Detection of Thrombin in Blood Serum by Using an Aptamer-Based Sensor. Angewandte Chemie International Edition 2005, 44 (34), 5456–5459.

42. Shoorideh, K., Chui, C. O., On the origin of enhanced sensitivity in nanoscale FET-based biosensors. Proceedings of the National Academy of Sciences 2014, 111 (14), 5111.

43. Kesler, V., Murmann, B., Soh, H. T., Going beyond the Debye Length: Overcoming Charge Screening Limitations in Next-Generation Bioelectronic Sensors. ACS Nano 2020, 14 (12), 16194–16201.

44. Mannoor, M. S., James, T., Ivanov, D. V., Beadling, L., Braunlin, W., Nanogap Dielectric Spectroscopy for Aptamer-Based Protein Detection. Biophysical Journal 2010, 98 (4), 724–732.

45. Arroyo-Currás, N., Dauphin-Ducharme, P., Ortega, G., Ploense, K. L., Kippin, T. E., Plaxco, K. W., Subsecond-Resolved Molecular Measurements in the Living Body Using Chronoamperometrically Interrogated Aptamer-Based Sensors. ACS Sensors 2018, 3 (2), 360–366.

46. Wilson, B. D., Soh, H. T., Re-Evaluating the Conventional Wisdom about Binding Assays. Trends in Biochemical Sciences 2020, 45 (8), 639–649.

