## Supplementary material for "Accelerated electron transfer in nanostructured electrodes improves the sensitivity of electrochemical biosensors": Fu_Seo_Kesler_SI

K. Fu et al.

### Contents

Supplementary **Table S1 |** Summary of properties for planar and nanoporous gold electrodes produced under different conditions.

Supplementary **Table S2 |** Coefficients of variation (CV) of signal level for planar and nanoporous gold electrodes obtained from different target concentrations.

Supplementary **Figure S1 |** Representative SWV plots from nanoporous (left) and planar electrodes (right) comparing buffer-only signal with signal from 10 μM DOX.

Supplementary **Figure S2 |** Representative cyclic voltammograms from nanoporous (red) and planar electrodes (blue), where the reduction peaks indicate the gold electrode surface area.

Supplementary **Figure S3 |** Aptamer probe density on nanoporous and planar electrodes after varying the incubated aptamer concentration from 100 nM to 10 μM.

Supplementary **Figure S4 |** Optimization of SWV parameters.

Supplementary **Figure S5 |** Signal gain at different ionic strengths.

### SUPPLEMENTARY TABLES

| | Average pore size [nm] | Signal gain at DOX 100 µM [%] |
|---|---|---|
| **Planar electrode** | **Non-porous** | **32.3 ± 14.5** |
| **No post-treatment** | **9.3 ± 3.6** | **193.8 ± 5.1** |
| **Thermal annealing only** | **24.1 ± 10.1** | **159.7 ± 16.1** |
| **Electrochemical coarsening only** | **33.7 ± 11.6** | **155.0 ± 21.3** |
| **Thermal annealing with electrochemical coarsening** | **63.1 ± 34.1** | **145.0 ± 22.2** |

**Table S1**: Summary of properties for planar and nanoporous gold electrodes produced under different conditions.

| **Target concentration [μM]** | **Nanoporous CV[%]** | **Planar CV[%]** |
|---|---|---|
| **0.1** | **3.85** | **19.59** |
| **0.3** | **3.87** | **19.56** |
| **1.0** | **3.47** | **19.72** |
| **3.0** | **3.37** | **19.82** |
| **10.0** | **2.47** | **19.53** |
| **30.0** | **3.84** | **18.84** |
| **100.0** | **5.21** | **18.30** |

**Table S2**: Coefficients of variation (CV) of signal level for planar and nanoporous gold electrodes obtained from different target concentrations. Data are averaged from six replicates (n = 6).

**SUPPLEMENTARY FIGURES**

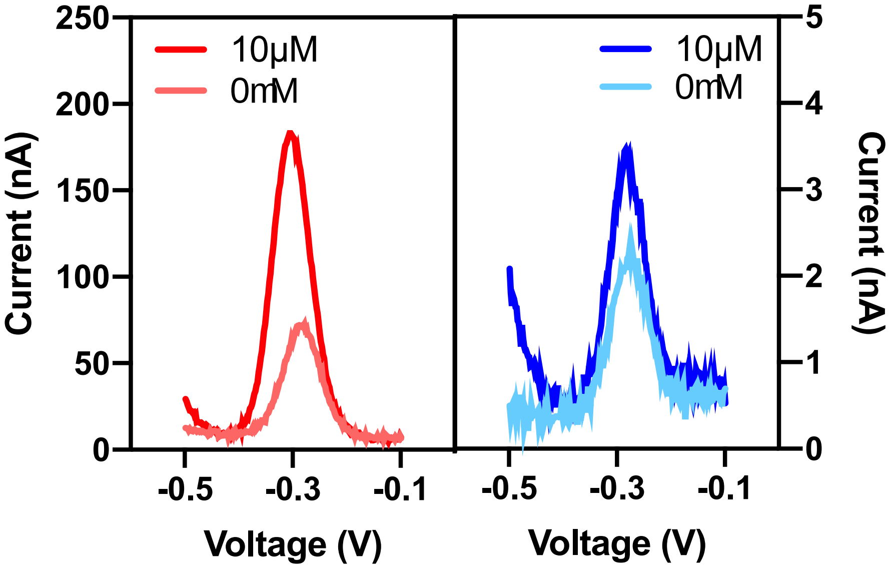


**Figure S1.** Representative SWV plots from nanoporous (left) and planar electrodes (right) comparing buffer-only signal with signal from 10 μM DOX.

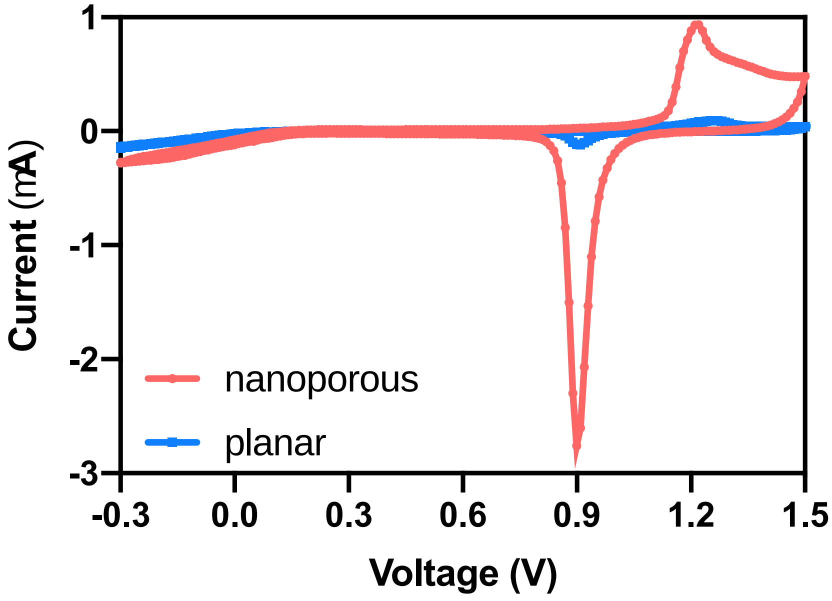


**Figure S2.** Representative cyclic voltammograms from nanoporous (red) and planar electrodes (blue), where the reduction peaks indicate the gold electrode surface area.

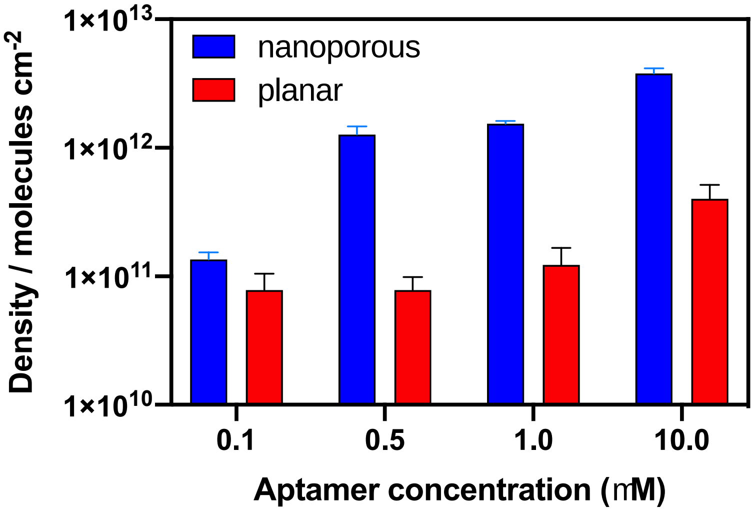


**Figure S3.** Aptamer probe density on nanoporous and planar electrodes after varying the incubated aptamer concentration from 100 nM to 10 μM. Datapoints are averages from seven replicates (n = 7).

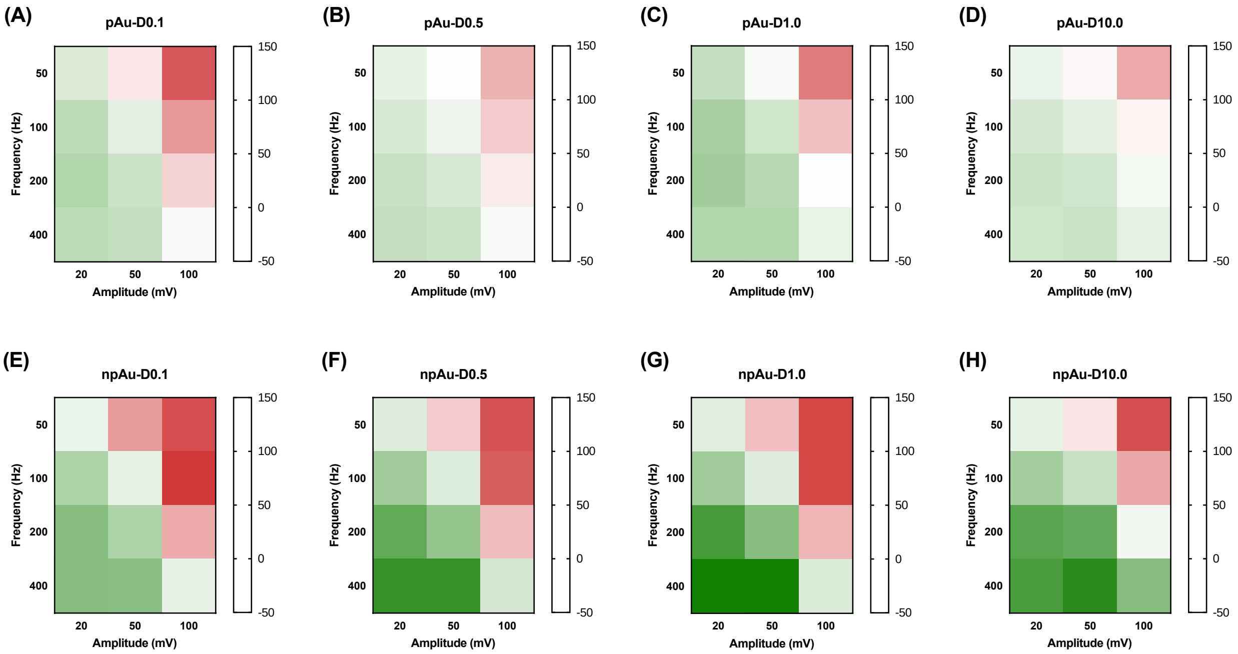


**Figure S4.** Optimization of SWV parameters. Heat-map of signal gain of electrochemical sensors with nanoporous (**A** to **D**) or planar electrodes (**E** to **H**) at different frequencies (50–400 Hz) and amplitudes (20–100 mV). Panels from left to right represent a shift from low to high aptamer density. All data are averaged over six replicates (n = 6).

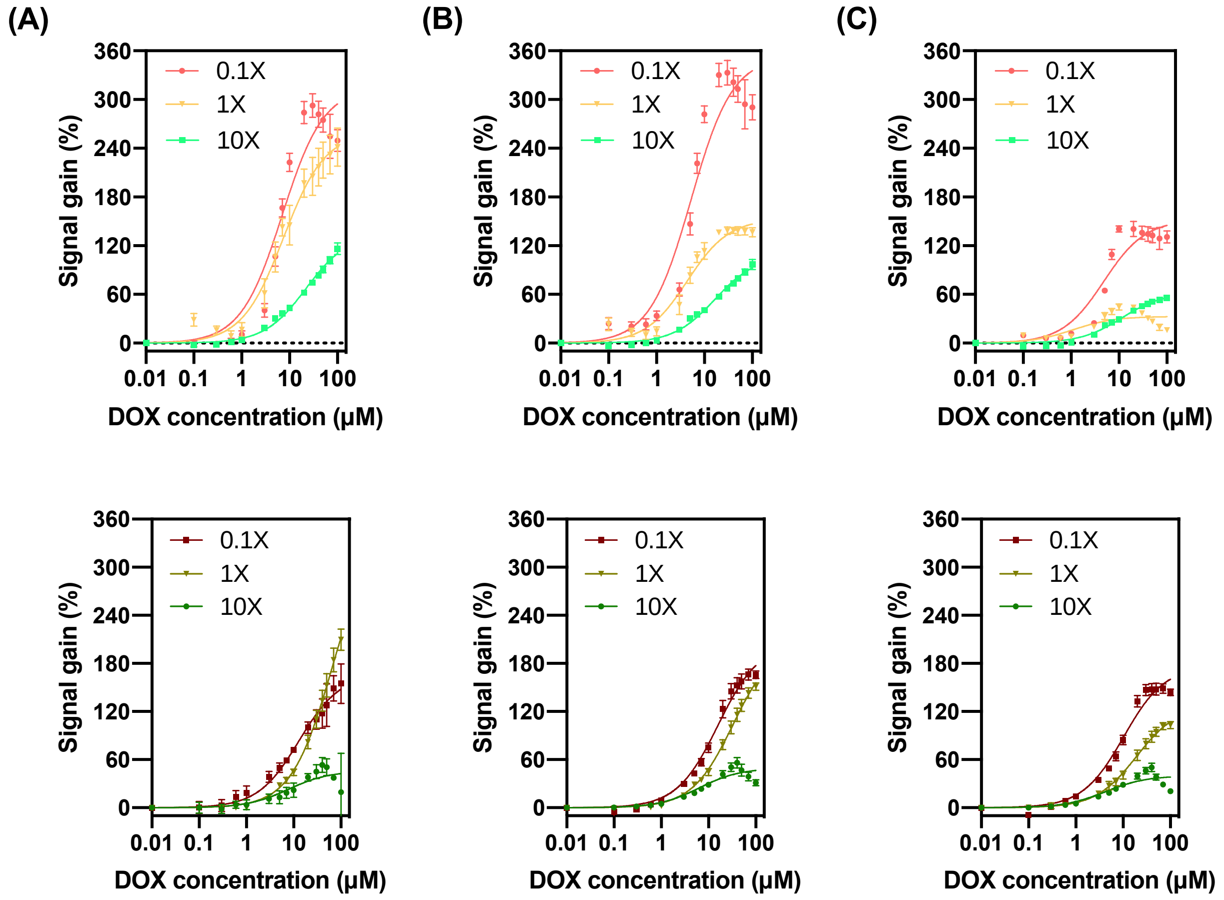


**Figure S5**. Signal gain at different ionic strengths. Signal gain is shown from nanoporous (top) and planar (bottom) electrodes in 0.1X, 1X and 10X SSC buffer at three SWV frequencies: (**A**) 400 Hz, (**B**) 200 Hz, and (**C**) 100 Hz. Datapoints are averaged over three replicates (n = 3). Notably, the effects of changing ionic strength on SWV were visible at lower frequencies on planar electrodes than on nanoporous electrodes; at higher frequencies, the signal gain curves of both types of electrodes converged towards their maximum values regardless of ionic strength. We speculate that these differences are caused by the difference in morphology between the flat planar and curved nanoporous interfaces.
